# Cannabidiol Interactions with Voltage-Gated Sodium Channels

**DOI:** 10.1101/2020.06.15.151720

**Authors:** Lily Goodyer Sait, Altin Sula, David Hollingworth, Benjamin J. Whalley, Rohini R. Rana, B.A. Wallace

## Abstract

Voltage-gated sodium channels are targets for a range of pharmaceutical drugs developed for treatment of neurological diseases. Cannabidiol (CBD), the non-psychoactive compound isolated from cannabis plants, was recently approved for treatment of two types of epilepsy associated with sodium channel mutations. This study used high resolution X-ray crystallography to demonstrate the detailed nature of the interactions between CBD and the NavMs voltage-gated sodium channel, showing CBD binds at a novel site at the interface of the fenestrations and the central hydrophobic cavity of the channel. Binding at this site blocks the transmembrane-spanning sodium ion translocation pathway, providing a molecular mechanism for channel inhibition. Modelling studies illuminate why the closely-related psychoactive compound THC may not bind to these channels. Finally, comparisons are made with the TRPV2 channel, also recently proposed as a target site for CBD. In summary, this study provides novel insight into a possible mechanism for CBD with sodium channels.

## Introduction

Voltage gated sodium channels (Navs) specifically enable the passage of sodium ions across cell membranes, contributing to the electrical signalling in cells (Ahern *et al*, 2016). The nine homologous mammalian sodium channel subtypes, designated hNav1.1-hNav1.9 (Supplementary Figs. 1 & 2), have different functional characteristics and expression profiles within different tissues (Catterall et al, 2005). Mutations of hNavs have been associated with a range of channelopathies, including pain, epilepsy, and heart disorders, making them major targets for drug development (Bagal et al, 2015, Kaplan et al, 2016).

Cannabinoids (including cannabidiol (CBD) and tetrahydrocannabinol (THC)) are hydrophobic compounds (Supplementary Fig. 3) produced by the cannabis plant. Whilst THC has been primarily associated with psychoactive drug use (Rosenberg *et al*, 2015; Pisanti et al, 2017), the non-psychoactive component, CBD, has been extensively investigated for its potential clinical applications as a therapeutic drug for treatment of epileptic conditions (Rosenberg et al, 2017). One formulation of CBD (GW Research Ltd., UK) has recently been approved by the European Medicines Agency and the Federal Drug Administration for use in children for the treatment-resistant epilepsies Dravet Syndrome and Lenox-Gastaut Syndrome (Sarker & Nahar, 2020). Both of these diseases are rare early onset epilepsies associated with Navs, with Dravet patients often having mutations in the hNav1.1 human sodium channel gene *SCN1A* (Marini *et al*, 2011). Furthermore, CBD has been shown to attenuate seizures and social deficits in a mouse model of Dravet syndrome (Kaplan et al, 2017). Despite a significant amount of evidence reporting on the effectiveness of CBD for treating epileptic conditions (Cross *et al*, 2017; Devinsky *et al*, 2018), the molecular basis of its target interactions remain unclear (Watkins, 2019).

Electrophysiology studies, however, have indicated that CBD can modify sodium channel functioning: at 10 μM, it significantly reduced action potentials in rat CA1 hippocampal neurons, as well as the Nav current density in human blastoma cells and mouse cortical neurons (Hill *et al*, 2014). In addition, CBD has been shown to inhibit the channel activities of human Nav1.1 to Nav1.7 isoforms, as well as those of the prokaryotic Nav homologue NachBac, with IC50s ranging from 1.5 to 3.8 μM, which suggests inhibition at physiologically-relevant concentrations. Functional studies (Patel *et al*, 2016; Ghovanloo *et al*, 2018) on both hNavs and the homologous prokaryotic Navs suggested CBD interferes with the inactivation processes of these channels. In a recent study (Mason & Cummins, 2020) CBD was shown to inhibit both resurgent and persistent sodium currents of hNav1.2 at concentrations of 1 μM. However, to date, the structural basis of the interactions of CBD and sodium channels have not been identified on a molecular level.

CBD has also been suggested to be a potential inhibitor of the Transient Receptor Potential Cation Channel Subfamily V Member 2 (TRPV2) channel (Qin *et al*, 2008; Morelli et al, 2013), which facilitates the non-specific movement of both sodium and calcium ions through plasma membranes. According to electrophysiology studies, CBD activates rat TRPV2 with an EC_50_ of 3.7μM (Qin *et al*, 2008), although the link with epilepsy (Morelli *et al*, 2013) is much less direct than that for sodium channels. Recently cryo-electron microscopy (cryo-EM) was used to elucidate the structure of TRPV2 in a CBD-bound state at a nominal resolution of 3.2 Å (Pumroy *et al*, 2019); that study indicated the presence of CBD in the pore region of the protein structure, thus supporting the proposal for TRPV2 being a candidate target for CBD binding.

In the present study, in order to examine the nature of the interactions of CBD with sodium channels, the high-resolution crystal structure of a complex of CBD with the NavMs voltage-gated sodium channel from *M. marinus* (Sula *et al*, 2017), was determined, enabling the binding sites for the CBD molecule to be clearly defined at high resolution. The NavMs channel has been shown to be a good exemplar for hNavs as they exhibit not only functional (Bagneris *et al*, 2014; Ulmschneider et al, 2013; Ke *et al*, 2018), but also sequence and structural homologies (Supplementary Figs. 1 and 2) (Sula *et al*, 2017; Sula & Wallace, 2017). The NavMs-CBD structure described in this work hasenabled comparisons of that binding site with binding sites of other sodium channel ligands and modifiers in human sodium channels, including the hNav1.1 and hNav1.2 channel, which are the predominant hNav isoforms found in human brain tissue. It also provides a means for comparing its binding site for CBD with that found in the TRPV2 channel.

This high resolution crystallographic study of a sodium channel–CBD complex thus provides a means of both understanding the molecular interactions of CBD and sodium channel targets, and how these may be related to its use for treatment of epilepsy.

## Materials and Methods

### Materials

Thrombin was purchased from Novagen Inc (Germany), decanoyl-N-hydroxyethylglucamide (Hega10) was purchased from Anatrace (USA), and dimethyl sulfoxide (DMSO), sodium chloride, 2-amino-2-(hydroxymethyl)-1,3-propanediol (Tris), and imidazole were purchased from ThermoFisher Scientific (USA). Purification columns were purchased from GE Healthcare (USA). Cannabidiol (CBD) samples were supplied by GW Research Ltd. (UK). The F208L (NavMsL) mutation was introduced using the SLIM site-directed mutagenesis protocol (Chiu *et al*, 2004), using the forward primer 5′-CTCACCACCCTGACCGTGCTCAACCTGTTTATTGG-3′ and reverse primer 5′-GAGCACGGTCAGGGTGGTGAGCATGATGAACGGGATG-3′. The sequence was verified by Source Bioscience, UK.

### Protein expression and purification

The NavMs (Uniprot ID A0L5S6) and NavMs_L_ proteins were expressed and purified as previously described (Sula *et al*, 2017), with the following modifications: the bound protein was eluted in a buffer containing 20 mM Tris, pH 7.5, 300 mM NaCl, 0.5 M imidazole and 0.52% Hega10. The Histag was removed by thrombin cleavage overnight at 4° C. The protein sample was loaded onto a Superdex 200 column and eluted with 20 mM Tris, pH 7.5, 300 mM NaCl, and 0.52% Hega10 buffer. Protein samples were pooled and concentrated to 10 mg/ml using a 100 kDa cut-off Amicon concentrator and stored at a concentration of 10 mg/ml at −80 °C.

### Crystallisation, Data Collection and Structure Determination

1 μl of cannabidiol (100 mM) in 100% DMSO was added to 50 μl of the purified protein solution to produce a final protein concentration of ~10 mg/ml containing 2 mM CBD and 2.5% v/v DMSO. The best crystals were grown at 4 °C via the sitting drop vapour diffusion method using a 2:1 ratio of the protein and reservoir solutions containing 0.1 M lithium sulphate, 0.1 M HEPES, pH 7, and 40% v/v PEG200. The apo NavMs_L_ crystals were grown under the same condition as the crystals of the CBD complex, but without the DMSO and drug. Crystals were flash-frozen, with the PEG200 acting as the cryo-protectant.

Data were collected on beamline P13 at the Electron Synchrotron (DESY, Germany); on beamline Proxima1 at the Soleil Synchrotron (France), and on beamlines IO3, IO4, and I24 at the Diamond Light Source (UK). Hundreds of crystals were screened and full data sets were collected from more than 40 crystals. Diffraction images were integrated and scaled using XDS (Kabsch, 2010) and then merged with Aimless (Evans & Murshudov, 2013) using the CCP4 suite of programmes (Winn *et al*, 2011). The structure was determined from the crystals which diffracted to the highest resolution (2.2 Å for the apo protein, and 2.25 Å for the CBD complex). Because of the small but significant variations in the unit cell dimensions and resolution between different crystals of the same type produced under the same conditions, as we have seen previously (Naylor *et al*, 2016; Sula *et al*, 2017), datasets from different crystals were not merged.

The structure determinations by molecular replacement were as previously described (Sula *et al*, 2017) using Phaser (McCoy *et al*, 2007) with the full-length wildtype NavMs structure (PDB 5HVX) as the search model. Model building was carried out using Coot (Emsley *et al*, 2010). Refinement was done using REFMACS (Murshudov et al, 2011). Data collection, processing and refinement statistics for both the apo and CBD complex structures are listed in Supplementary Table 1. The structure quality was checked using PROCHECK (Laskowski *et al*, 1993) and MolProbity (Chen *et al*, 2010), which indicated that 99.2% of the residues were in allowed conformations. Figures were created in CCP4mg (McNicholas *et al*, 2011), unless otherwise noted.

## Results

This study utilised the prokaryotic NavMs voltage-gated sodium channel, which has been previously shown to be an excellent structural and functional exemplar for human sodium channels, to examine the site of interactions of the naturally-occurring non-psychoactive CBD compound isolated from cannabis plants. One formulation of CBD (GW Research Ltd., UK) has recently been approved by the European Medicines Agency (EMA) and the Food and Drug Administration (FDA) (Sarker & Nahar, 2020) for treatment of specific and severe epilepsies. NavMs was not only chosen for this study because it exhibits both strong sequence and structural homology to hNavs (Supplementary Figs. 1 & 2), but it has also been shown to have highly similar functional (Bagneris *et al*, 2014), conductance (Ulmschneider et al, 2013), and drug binding (similar IC_50_ values) characteristics (Bagneris *et al*, 2014) as human Nav1.1 sodium channels. Whilst NavMs channels are tetramers with each monomer consisting of 6 transmembrane (TM) helices (4 of which form each of the voltage sensor subdomains and 2 of which form the pore subdomains), all of the hNav channel isoforms are monomers of four similar but not identical domains (each of which consists of 6 transmembrane helices that are comprised of 4-helical voltage sensor subdomains and 2-helical pore subdomains). The major difference between NavMs and the hNavs is the presence of the inter-domain loop regions in the human channels, (which also differ considerably between hNavs) (Supplementary Fig. 1, Fig. 5).

The major reason for using crystal structures of the NavMs channel for this study is that they provide the, to date, highest resolution (~2.2-2.5 Å) views of any sodium channel (Naylor et al, 2016; Sula et al, 2017), especially of the TM and drug binding regions, thus enabling detailed views of the protein molecular structures with drugs bound to them. In contrast, the cryo-EM structures of hNavs available to date generally have overall resolutions of between ~4-5 Å, with the transmembrane regions having the best resolutions of ~3 Å, with their extra- and intra-membranous regions being less well defined. However, these similarities would not be sufficient to indicate the value of using NavMs for understanding the molecular basis of drug interactions if their functional roles (conductance and drug binding affinities) were not comparable to those of hNavs. NavMs and hNav1.1 exhibit similar ion flux and conductance properties, and as well as very similar binding affinities for a wide range of sodium channel-specific drugs (Bagneris et al, 2014).

Structure/function/drug-binding studies using some of the other prokaryotic sodium channels have also showed their comparability to hNavs for drug binding (Payandeh et al, 2011; Jiang et al, 2019), and they too have been used for drug discovery projects (Martin & Corry, 2014; Ouyang et al, 2007), although their structures tend to be of lower resolution than those of NavMs. Much of the focus of this study has been on comparisons of NavMs with hNav1.2 and hNav1.1, as these are the sodium channels primarily found in human central nervous system tissues. Although as yet there is no structure available for the hNav1.1 channel, its strong sequence homology to the hNav1.2 has enabled the production of the molecular model used in the comparison studies described her Fig. 5).

### The CBD Binding Site in Sodium Channels

The CBD binding site is located and clearly visible in a well-defined region of the NavMs-CBD structure (Figs. 1A & B). It is sited in a hydrophobic pocket present in each subunit that runs perpendicular to the channel direction (Montini et al, 2018) (such features have been designated “fenestrations” and are located (horizontally in Fig. 1C) in the TM region, just below the level of the selectivity filter, and are the features originally proposed by Hille (1977) as sites for ingress of hydrophobic drugs into the channel interior. CBD is located at the end of the fenestration that lies closest to the central pore, and protrudes into (and blocks) the central transmembrane cavity, just below the sodium ion selectivity filter (Fig. 1C). There is enough room for four CBD molecules in this region, although one would be sufficient to block sodium ion passage, as seen from the HOLE (Smart *et al*, 1993) depictions (Fig. 2A and Supplementary Fig. 4) and pore radius plots (Fig. 2B), which show the size of the transmembrane pathway with and without different numbers of CBD; the blockage clearly provides a mechanism for channel inhibition as well as a basis for understanding the concentration-dependence of the drug effects. Each binding site is comprised of 11 residues from three different subunits in NavMs (shown in different colours in Fig. 3). The corresponding residues in hNavs are from three different domains of the same polypeptide chain (Figs. 4 & 5, Supplementary Fig. 5).

**Fig. 1.**
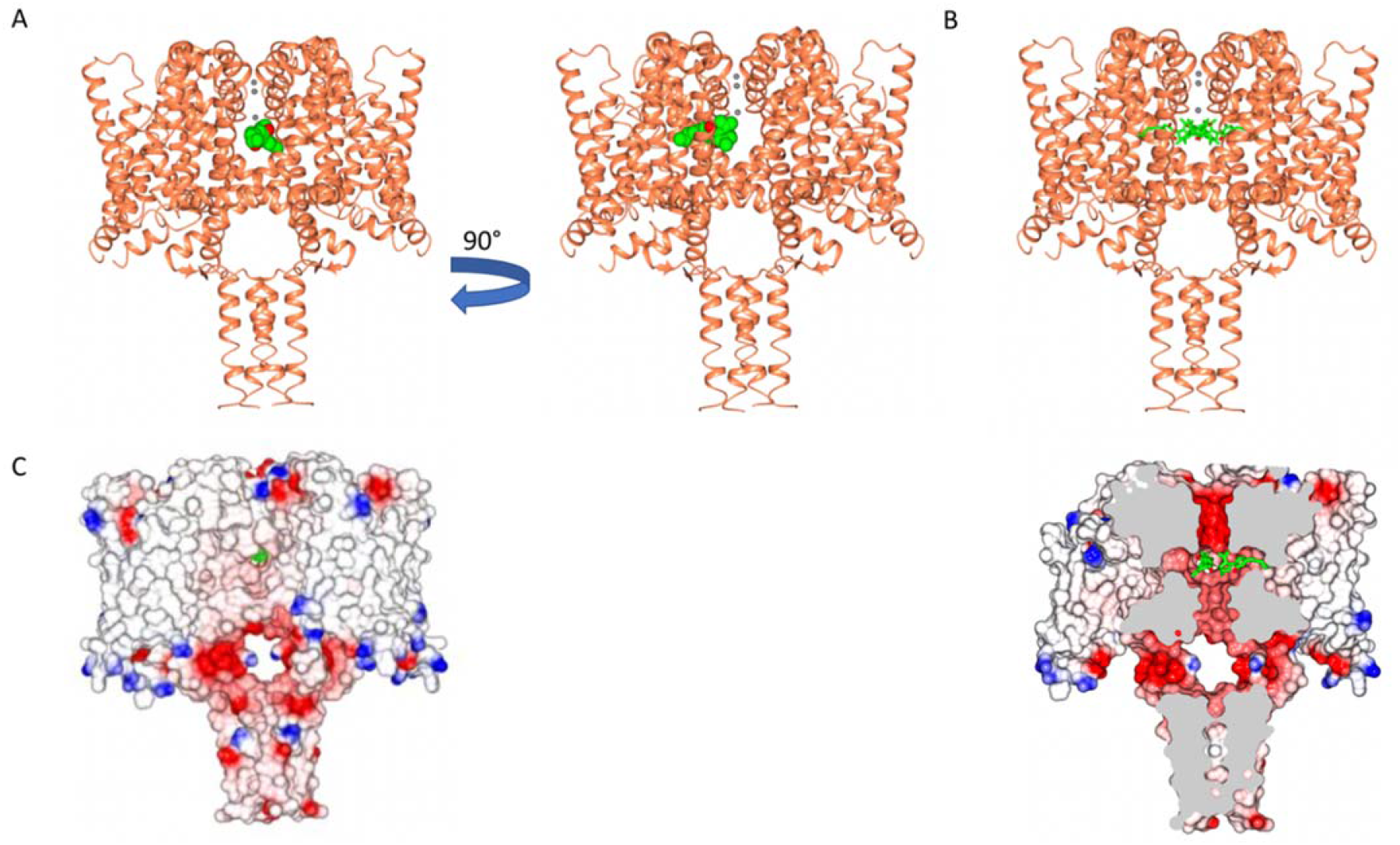
The NavMs Sodium Channel/Cannabidiol (CBD) Crystal Structure. A) The crystal structure (2.25 Å resolution) of the NavMs sodium channel (in coral coloured ribbon depiction) with one CBD molecule (in green space-filing depiction), showing its location within the hydrophobic cavity of the channels located within the fenestration. Three sodium ions are shown as grey spheres in the selectivity filter. The view on the right side is rotated 90 degrees from the view on the left. B) As in a) but showing 4 CBD molecules in the stick depiction) C) (left) Surface view of space filling structure of NavMs with CBD (in green) present. The orientation is the same as in the left panel of part A. The CBD is just visible through the fenestration hole. (right) As in left panel, but sliced through the space filling model (and through the middle of the fenestration) with 2 CBD molecules present, showing where the drug lies along the fenestration and ion pathway.

**Fig. 2.**
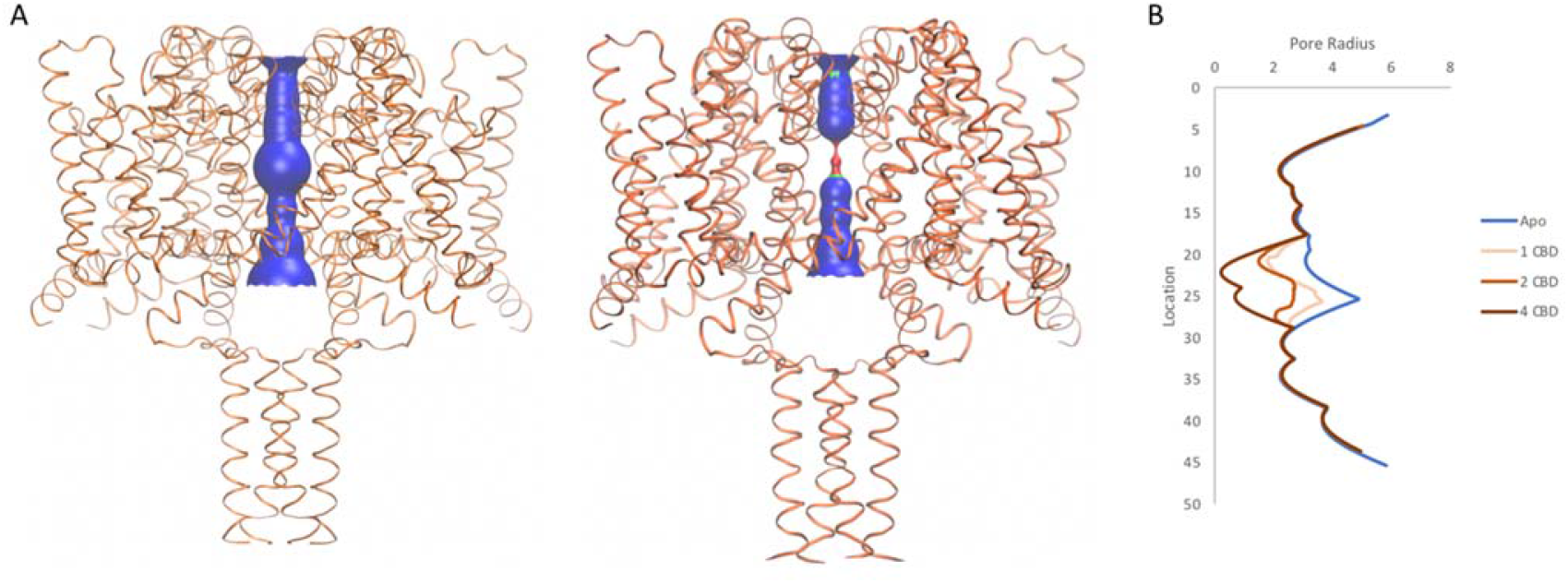
Pore Diameters in the Absence and Presence of CBD. A) The NavMs structure (coral) is depicted in ribbon motif. The diameter of the central transmembrane pore was calculated using the HOLE algorithm (Smart *et al*, 1993). The pore interior dimensions are shown for the (left) apo structure and (right) for the structure with 4 CBD molecules present. In this figure the HOLE surface is depicted in blue for pore radii greater than 2.3 Å, green for radii between 2.3 and 2.0, and red for radii less than 1.15 Å. There is no occlusion in the absence of CBD, so hydrated sodium ions could freely pass through the pore. The NavMs structure with 4 CBD molecules present shows a full occlusion in the middle of the transmembrane pathway (near the center of the hydrophobic cavity) of the channel, so ion transport would be prevented (Naylor *et al*, 2016). B) Accessibility plots of pore radii versus position in the pore, in the absence and presence of different numbers of CBD molecules. The plot for the apo structure is in blue, and the plots for the CBD-containing structures with 1, 2, or 4 CBD molecules are in coral, red and black, respectively. The plots for 2 and 3 CBD molecules were the same, so the latter is not shown. In cases where at least some region of the radius is <2.0 Å, sodium ions will not be able to be translocated across the channel (Naylor *et al*, 2016). This plot shows, therefore, that regardless of whether there 1 or more CBD molecules present, ion passage will not occur. These figures were produced using VMD software (Humphrey *et al*, 1996).

**Fig. 3.**
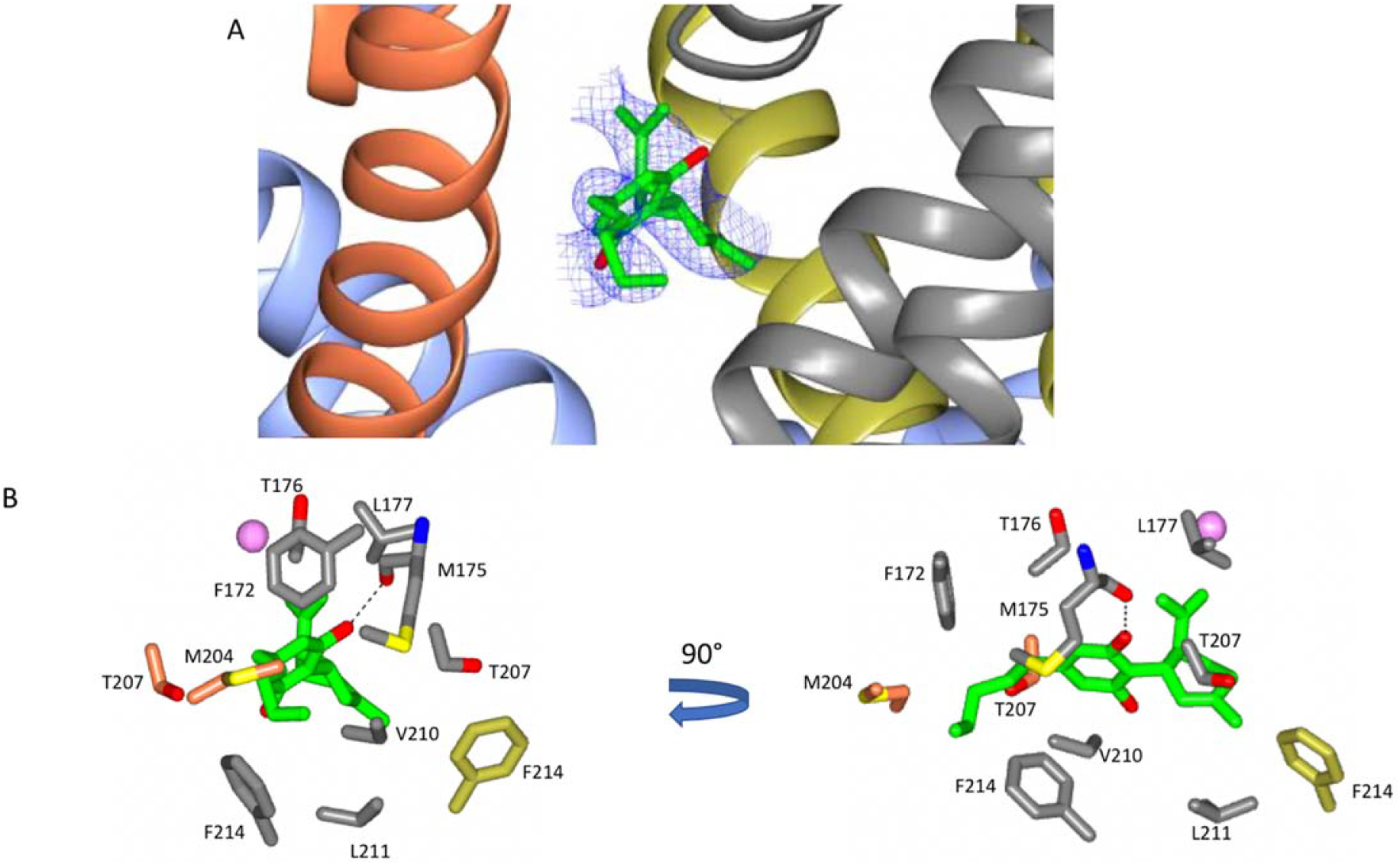
Detailed View of the CBD Binding Site. A) The polypeptide backbone of the NavMs-CBD complex is depicted in ribbon motif. The ribbons are coloured by subunit (only regions of the three subunits that come in close contact with the CBD, depicted in red, grey and yellow) are shown. The (2Fo-Fc) map (shown in blue mesh) was calculated at 1 sigma and the structure of the CBD molecule present is shown in stick depiction. B) (left) Detailed structures of residues that lie within 3.9 Å of the CBD molecule (which is depicted in green/red stick representation) are shown and coloured by domain (as in part A). The sphere indicates the sodium ion site furthest from the extra-membranous surface (the one located farthest into the channel). (right) The same view, rotated by 90 degrees.

**Fig. 4.**
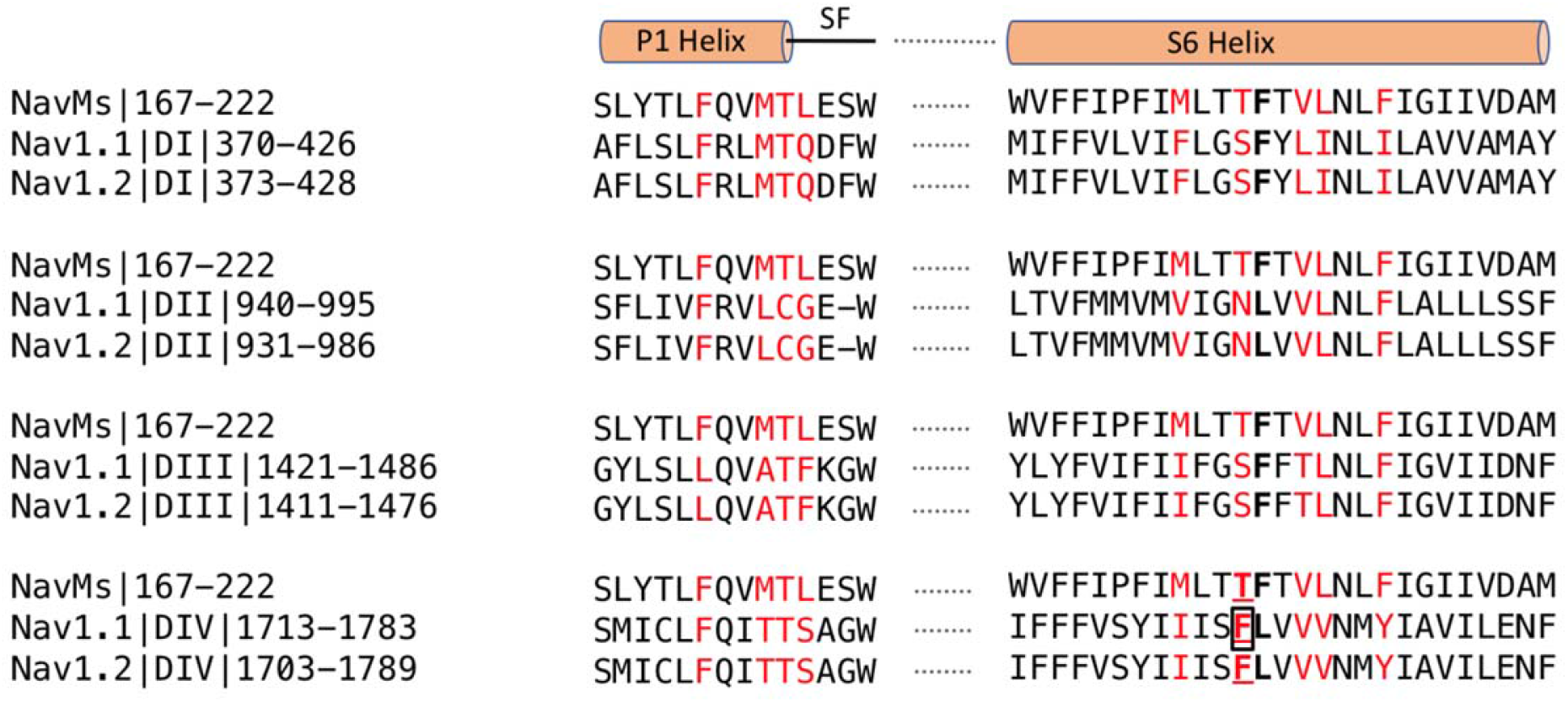
Sequence Alignments of CBD Binding Sites in NavMs, hNav1.1 and hNav1.2. In red are the CBD binding residues (with 3.9 Å of the compound) in NavMs and the equivalent residues in hNav1.1 and hNav1.2. The bold black F indicates the site of the NavMs F208L mutant used in these studies. It was changed from F to L in NavMs_L_ because in half of the human Nav domains it is an F and in the other half it is an L. However, as seen in Supplementary Fig. 8, the residue type present at this site makes essentially no difference in the structure. Binding residues occur within the P1 pore helix, the selectivity filter loop and the S6 helix. The residue, which when mutated to alanine in hNav1.1 reduces the binding affinity of CBD, is boxed. The sequence alignment was carried out using Clustal Omega (Siever et al., 2011) and annotated manually.

**Fig. 5.**
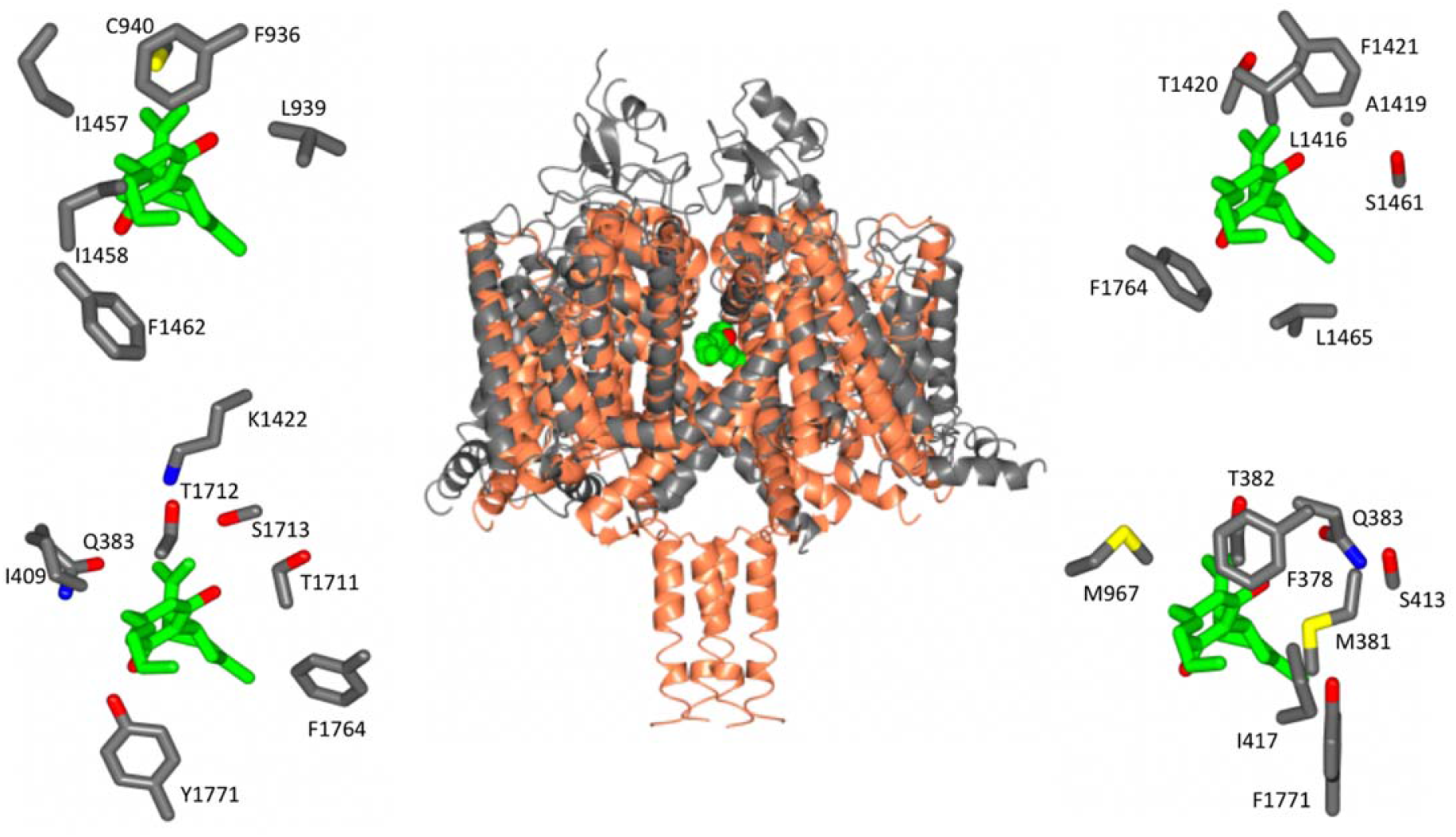
Location of Equivalent Binding Sites in NavMs and hNav1.2. (Middle) Structural alignment of the NavMs-CBD crystal structure (coral) and the hNav1.2 cryo-EM structure (grey). The RMSD of the aligned structures is 3.2 Å. (Top left): equivalent binding residues in domain 1 and domain 2 of hNav1.2 found within 4 Å of the CBD site. (Top right): binding residues in hNav1.2 between domains II and III located within 4 Å of the CBD binding site. (Bottom left): Residues in hNav1.2 between domains III and IV within 4 Å of the CBD binding site. (Bottom right): Residues in hNAv1.2 between domains IV and I within 4 Å of the CBD binding site. In the surrounding panels the atoms in the protein are coloured by atom type, with carbons represented in grey, oxygen in red, nitrogen in blue, and sulphur in yellow, whilst the carbon atoms of the drug are depicted in green.

The location of this binding site is very close to the locations of the binding sites that have been identified for analgesic and other hydrophobic compounds in both NavMs (Bagneris *et al*, 2014) (Supplementary Fig. 6) as well as in another bacterial sodium channel, NavAb (El-Din *et al*, 2018). This is of interest because those other compounds also inhibit hNav functions, and so suggest the importance of this site for drug interactions in humans. All of the interactions seen except one, that of residue M175 (Fig. 3B), involve hydrophobic interactions rather than hydrogen-bond formation (but that particular interaction between the main chain carbonyl of residues M175 and the OH group present in CBD, may be important for specificity of binding – see next section). It should be noted here, that this region of the apo structure also exhibits some electron density (but has a different size and shape) that has been attributed to detergent molecules, The (2Fo-Fc) electron density map of the CBD complex (Fig. 3A) clearly indicates that in these crystals the site is occupied by CBD rather than detergent.

### The Molecular Basis of CBD Binding Relative to THC

There are two main cannabinoids that can be extracted from cannabis plants, the psychoactive tetrahydrocannabinol (THC) and the non-psychoactive cannabinoid (CBD). Electrophysiological studies on hNavs and the bacterial NachBac have identified CBD (Ghovanloo *et al*, 2018, Patel *et al*, 2016) as having functional effects that are distinct from those of THC on these channels.

The chemical structures of CBD and THC are very similar (Supplementary Fig. 3), differing only by the presence of an additional free hydroxyl group on one of the rings in CBD (in THC the equivalent oxygen forms part of a closed pyran ring). Therefore, the structure of the CBD/NavMs complex was examined to see if it could provide a clue as to the reasons for the different functional effects of the two compounds. As can be seen in Supplementary Fig. 7, by placing the THC structure into the CBD binding site with the same orientation as found for CBD, it can be physically and sterically accommodated. However, and crucially, it is missing the one electrostatic interaction seen between CBD and NavMs: the hydrogen bond between the oxygen of the main chain residue M175 and the drug. This is the consequence of the absence of the additional free hydroxyl group in THC, as noted above. That hydroxyl group is the one which forms the hydrogen bond present in the CBD-protein complex. This provides an additional intermolecular interaction for CBD, and could account for the differences in binding affinities of the two compounds (Ghovanloo et al, 2018) [as well as (possibly) the differences in psychoactive properties of the compounds].

### Specificity/Potential Interactions with Other hNavs

The focus of functional effects of CBD on hNavs has primarily been on hNav1.1, due to its association with epilepsy, although there is yet no structure for this isoform. However, it has been possible to examine potential interactions using a hNav1.1 homology model based on the hNav1.2 cryo-EM structure (Supplementary Fig. 5), which suggests, not surprisingly, that the interactions would be very similar to those of Nav1.2. In NavMs, the involvement of T207 residue is of importance as it is well established as the primary binding site for local anaesthetics. Furthermore, when the equivalent residue (F1774) was mutated in hNav1.1 the binding affinity of CBD was found to decrease by a factor of 2 (Ghovanloo *et al*, 2018). The binding site residues (coloured red in Fig. 4) include both residues that are identical/homologous in NavMs and hNavs as well as residues that are only found in NavMs and not in human Navs. In most cases the non-cognate residues are also variable between hNavs and would thus appear not to be essential for the interactions.

### Comparison with Binding to the TrpV2 Channel

A recent study (Pumroy *et al*, 2019) has described the interaction of CBD with the TRPV2 channel, as demonstrated by a cryo-EM investigation of its complex, solved at an overall resolution of 3.2 Å. The location of the CBD was visible in the structure, as were the general features of the binding site. Although the lower resolution of that structure did not allow detailed analysis of its binding site, it was clear that involved a number of hydrophobic side chains, and required a partial refolding of the adjacent region of the protein polypeptide. The binding site found for CBD in the TRPV2 structure is in a similar region to that of CBD in NavMs (Fig. 6). However, the sodium channel CBD site is located further into the fenestration than it is in TRPV2, but closer to the ion binding sites and thus would more effectively block the transmembrane passageway for ion conductance.

**Fig. 6.**
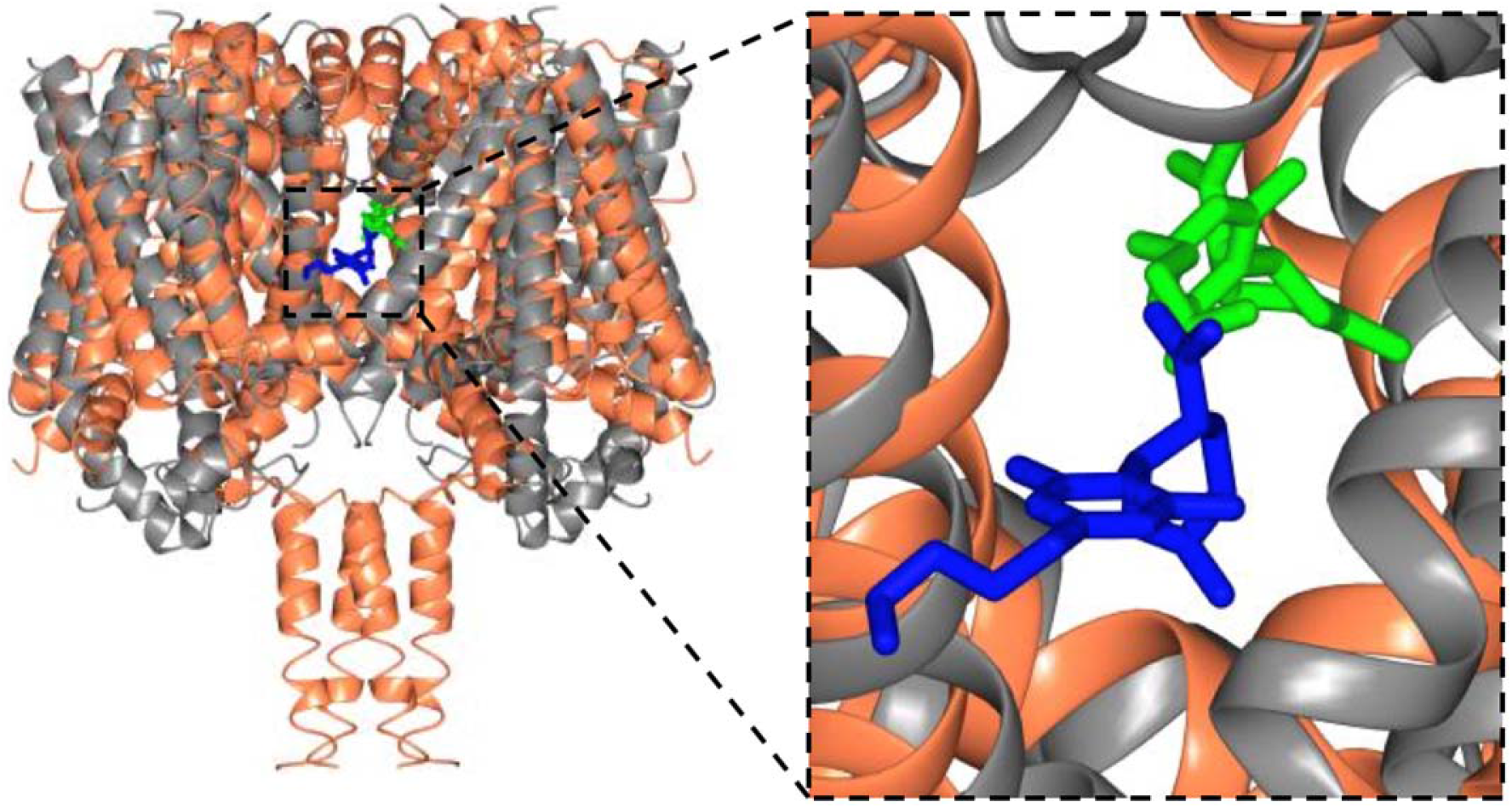
Structural Alignments Between the NavMs-CBD (coral ribbons) Crystal Structure and the TRPV2-CBD cryo-EM Structure (grey ribbons) (Left) Overall alignment of the structures. The TRPV2 structure was trimmed to remove the disordered regions for clarity. The RMSD of the alignment is 4.2 Å. The CBD in NavMs is in green and that in TRPV2 is in blue. CBD in NavMs appears to be located further into the fenestration than it is in TRPV2. (Right) Detailed view of the CBD sites, highlighting the similarity and differences in orientation and location in the two channel types.

## Conclusions

This study has demonstrated the nature of the interactions of CBD and a voltage-gated sodium channel, showing that CBD binding blocks the transmembrane pathway for sodium ion translocation through the membrane (Naylor et al, 2016), and hence provides a potential mechanism for the functioning of CBD in sodium channels. This further suggests a possible molecular basis for the medicinal effects of CBD in the treatment of epilepsies, as sodium channels have been shown to be causally-related to various types of human epilepsy, with disease-related mutations interfering with sodium ion transmembrane flux. The CBD binding site is a novel site, near to, but not coincident with, known analgesic binding sites in sodium channels; binding at this site would effectively block sodium channel functioning. The binding site is located at the pore end of the transmembrane fenestrations which enable the ingress of hydrophobic molecules into the channel lumen, hence indicating this may also provide the pathway for CBD to enter and block the channels.

Examination of the residues involved in the binding site interactions and modelling of the THC into the CBD binding site have indicated a possible reason for why the closely-related psychoactive cannabinoid THC, has not been observed to have a similar effect on epilepsy nor on sodium channel function: THC would be able to physically fit in the site when oriented in the same manner, but it does not have the same hydroxyl moiety that in CBD forms an important hydrogen-bonding interaction with the channel protein.

Recent cryo-EM structural studies (at lower resolution) have suggested that the TRPV2 channel may be the CBD binding target, although that study did not show the relationship of the binding site to epilepsy-based mutations. However, whilst the TRPV2 channel has a quite different overall fold from that of sodium channels and it acts as a conduit for much larger substrates, it is interesting that the binding site for CBD in TRPV2 appears to be in a roughly comparable structural feature near the transmembrane substrate pathway to that found in this study for the ion pathway in sodium channels.

In summary, this study has provided high resolution structural evidence for the basis of the molecular interactions of CBD, a drug recently approved for treatment of epilepsy, with a voltage-gated sodium channels target. The described structural work can therefore guide further functional studies to explore differential CBD selectivity for human Nav isotypes and their relevance to clinical studies, thus shedding further light on the polypharmacological profile of CBD.

## Author Contributions

BAW – Conceptulization; BAW, AS, LGS – designed experiments; LGS, AS, DH – purified and crystallised protein; AS, LGS, DH – collected crystal data, undertook structure solutions and analyses, produced Fig.s and tables; AS – did PDB depositions; BJW, RRR - valuable discussions that led to development of this research project, GW Research Ltd. – provision of a generous supply of CBD; BAW – original draft of paper and supervision of the project; all authors contributed to the writing of paper and approved the final draft.

## Funding acquisition

Grants BB/L006790 and BB/R001294 from the U.K. Biotechnology and Biological Science Research Council [BBSRC] (to BAW). Ph.D. studentship from the UCL-Birkbeck Medical Research Council DTP programme (to LGS). Beamtime grants for access to the DESY (Germany), Soleil (France) and Diamond (UK) synchrotrons (Birkbeck/UCL BAG consortium). The funding sources were not involved in the study design, data collection and interpretation, or decision to submit the work for publication.

## Datasets/Availability

Structure factors and coordinates for the apo and CBD-bound NavMs_L_ structures have been deposited in the Protein Data Bank under accession codes PDB6YZ62, and PDB6YZ0, respectively.

## Supplementary Material (Sait, Sula et al)

**Supplementary Fig. 1.**
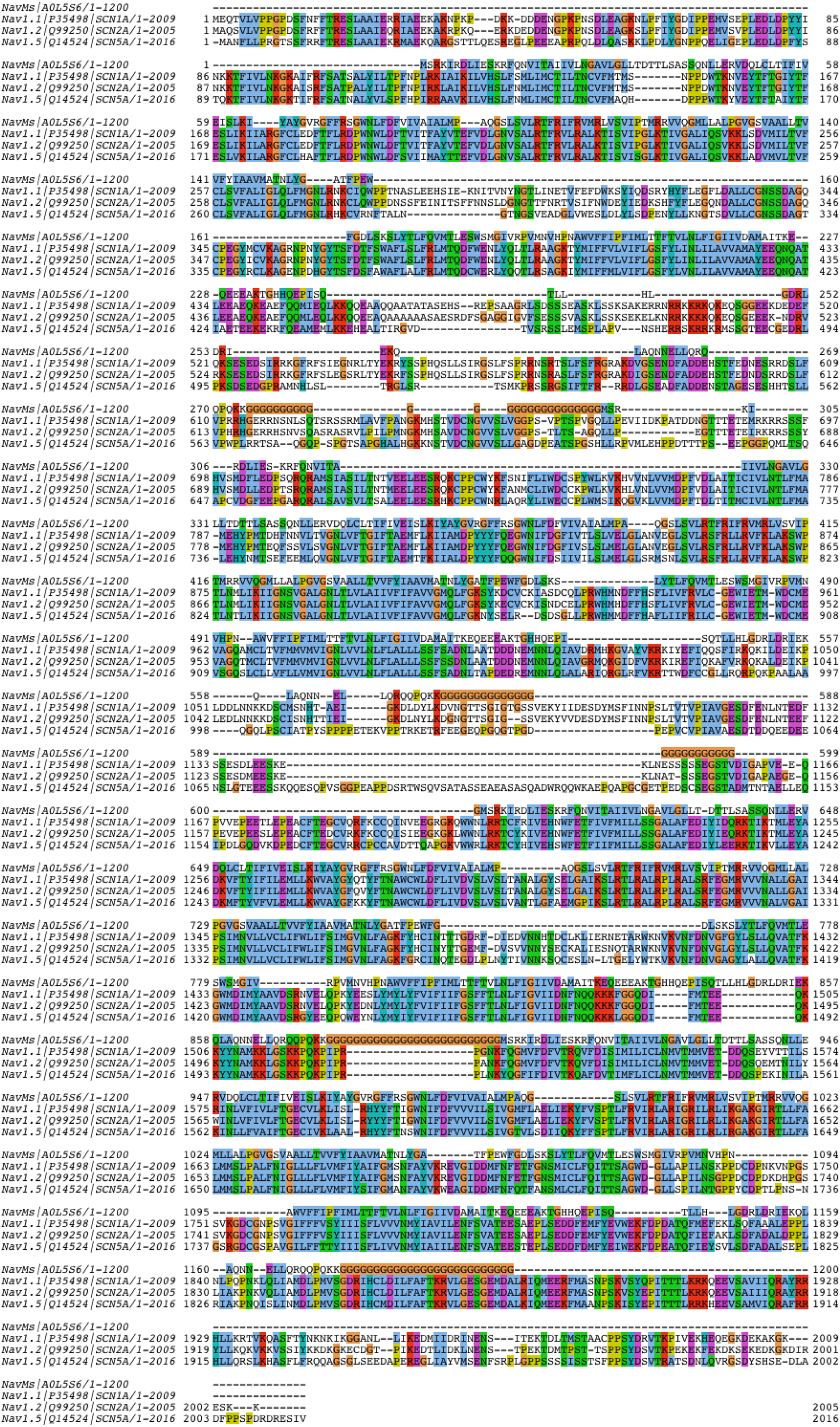
Alignment of Full Length Sequences of NavMs, hNav1.1, hNav1.2, and hNav1.5. This sequence alignment was carried out using Clustal Omega (Siever et al., 2011), visualized using Jalview (Waterhouse et al., 2009) and annotated using Clustal X coloring scheme (Waterhouse et al., 2009).

**Supplementary Fig. 2.**
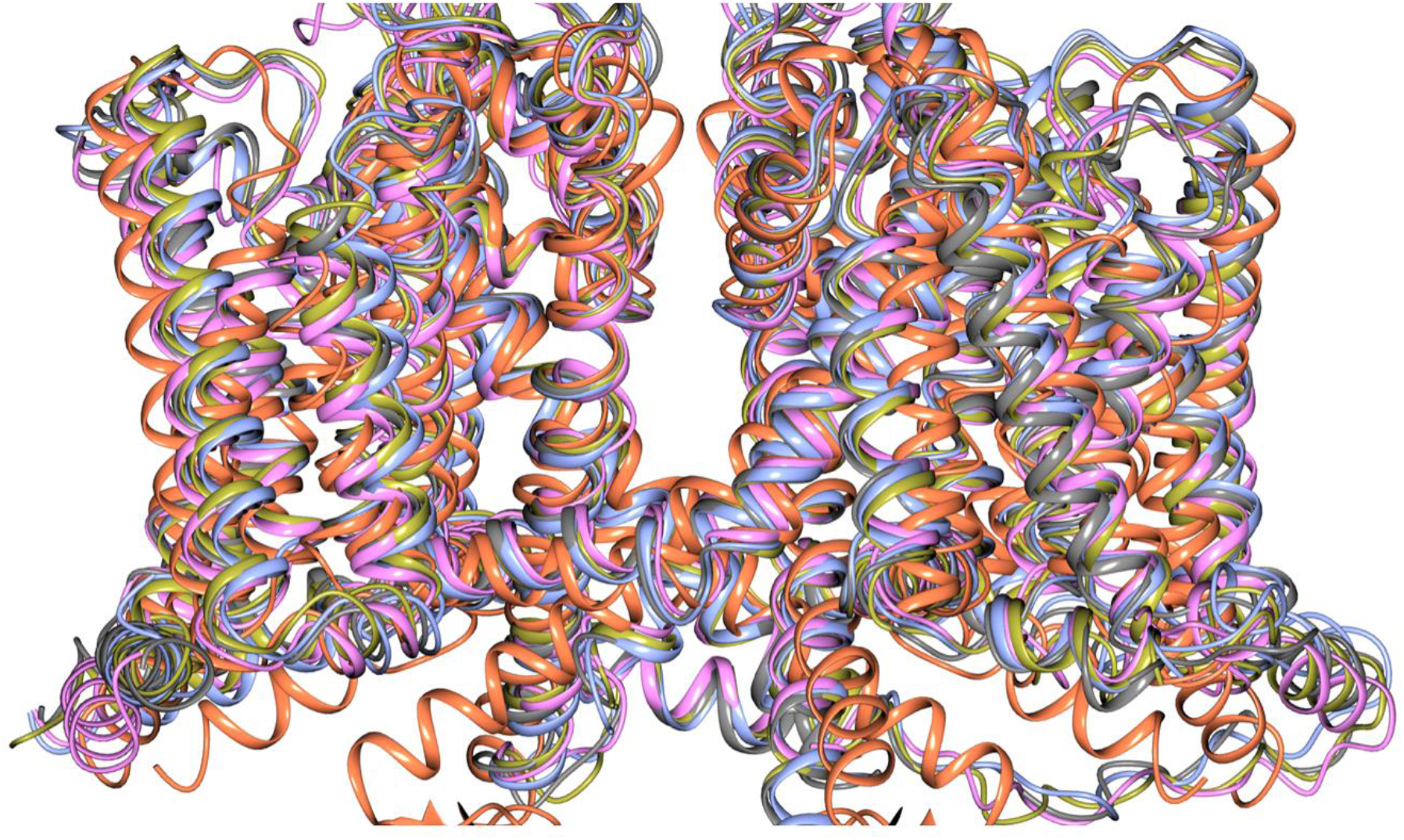
Overlay of Crystal Structure of NavMs (coral) and CryoEM Structures of hNavs 1.2 (grey), 1.4 (blue), 1.7 (gold) and rat Nav1.5 (pink).

**Supplementary Fig. 3.**
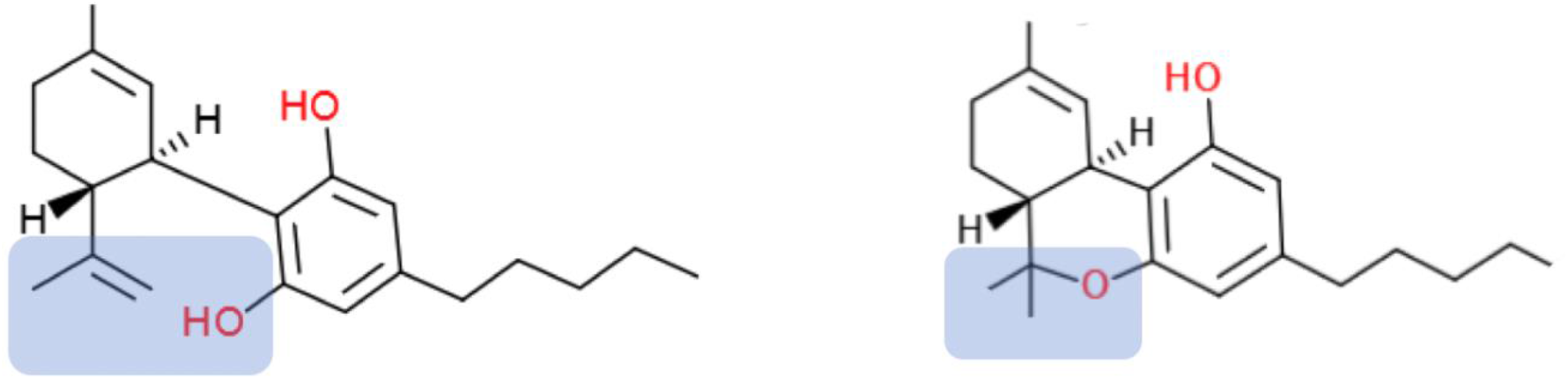
Chemical Structures of Cannabidiol (CBD) (left) and Tetrahydrocannabinol (THC) (right). The difference between the two structures is highlighted in blue background. The formation of pyran ring in THC removes the hydrogen from the hydroxyl group which is present in CBD and form a hydrogen bond in the channel-CBD complex.

**Supplementary Fig 4.**
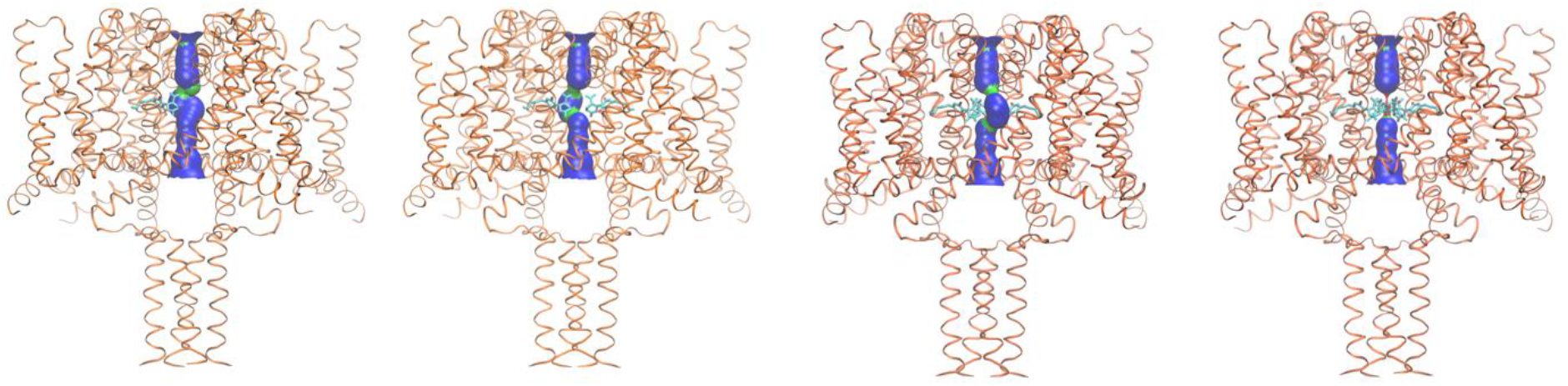
HOLE Depiction of the Dimensions NavMs-CBD Complexes with Different Numbers of CBD Molecules Present. This depiction is as in Fig. 2. The protein is depicted in coral ribbons, and the central pore size is indicted using the HOLE algorithm. Figures from left to right show the effects of increasing numbers (1, 2, 3 and 4) CBD molecules (depicted in stick representation) on the side of the central pore. The regions where the pore is sufficiently wide to enable the passage of sodium ions is in blue (pore diameter greater than 4.6 Å) surface representation, and regions where the pore is narrowed which might enable the passage of sodium ions is in green (pore diameter between 2.3 Å and 4.6 Å).

**Supplementary Fig. 5.**
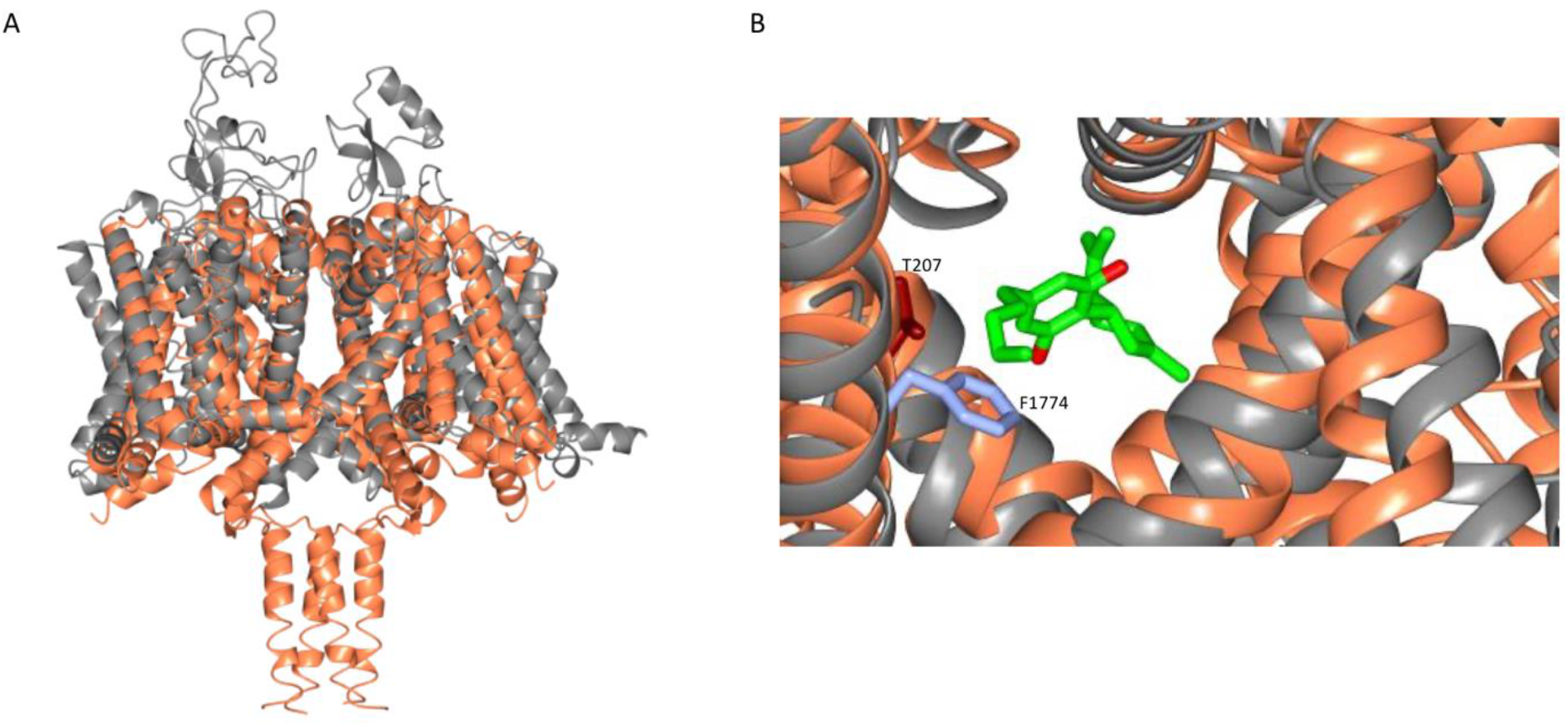
Alignment of the NavMs-CBD Structure (coral) with the Homology Model of Nav 1.1 (grey). The homology model was created in SWISS-MODEL (Waterhouse *et al*, 2018) using hNav1.2 (PDB code 6J8E) as a template. A) Overall structural alignment. (RMSD =3.34 Å). B) Detail of overlay in the region of residue F1774 (ice blue) in hNav1.2; this is a residue that has been identified as being in the local anesthetic binding site, and was the residue that was mutated in (Ghovanloo *et al*, 2018), which in their electrophysiology experiments showed effects on CBD inhibition. This residue coincides with the location of T207 (red) in NavMs.

**Supplementary Fig. 6.**
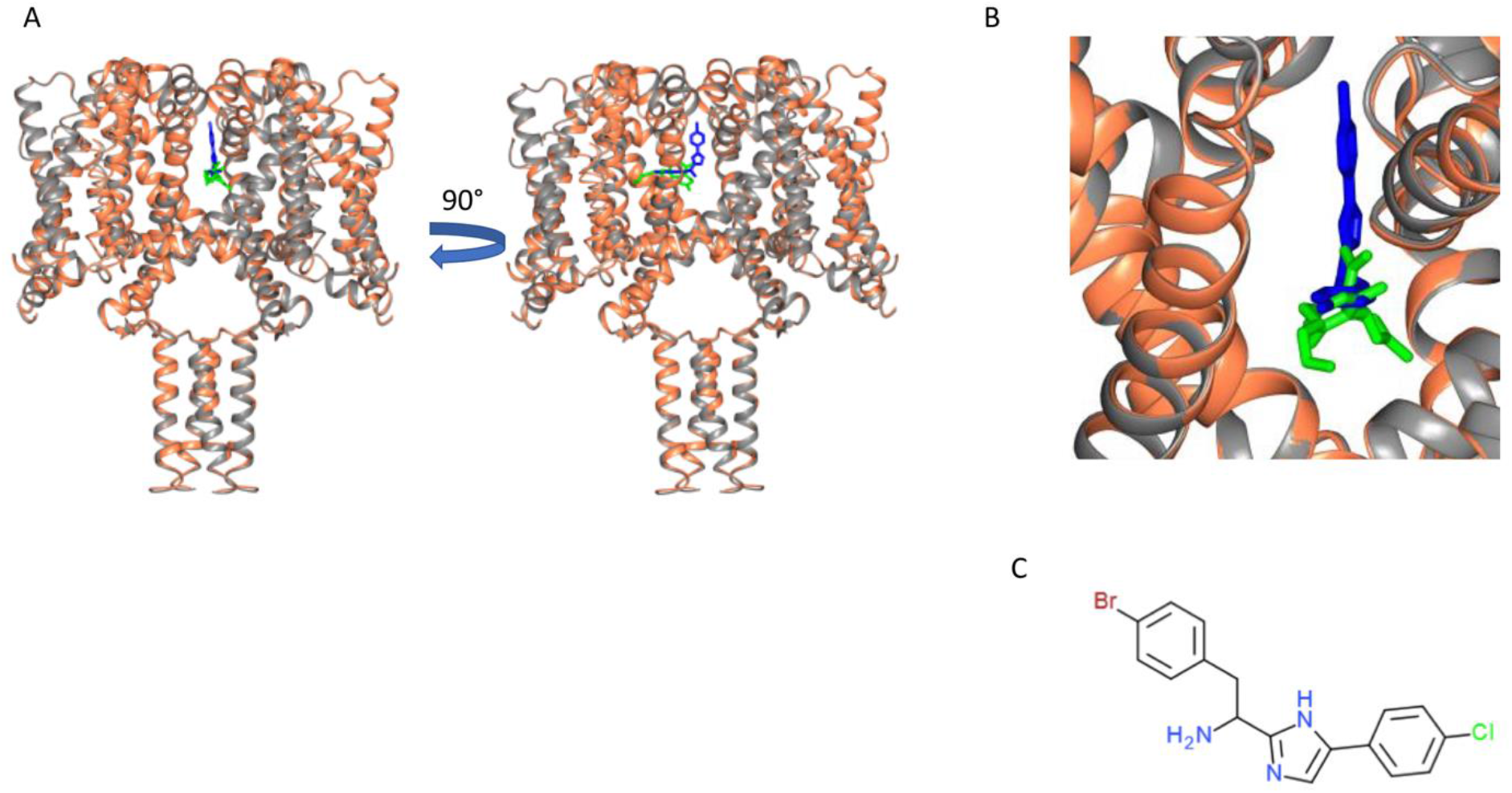
Similarity of CBD and Analgesic Compound Binding Sites. A) Structural alignment of NavMs_L_-CBD complex (protein in coral ribbon depiction, CBD in green stick depiction), with NavMspore-PI1 complex structure (Bagneris *et al*, 2014). The NavMs pore protein is in grey ribbon depiction and the PI1 molecule is in blue stick depiction. PI1 is a highly potent designed analgesic compound which binds to and inhibits flux through the NavMs channel (Bagneris *et al*, 2014). Two views of the aligned structural complexes are shown, rotated by 90 degrees, which show the similarity, but not identity of the binding sites of the two ligands. B) Detailed view showing the locations of these molecules in the pore/fenestration area. C) Chemical structure of the PI1 compound.

**Supplementary Fig. 7.**
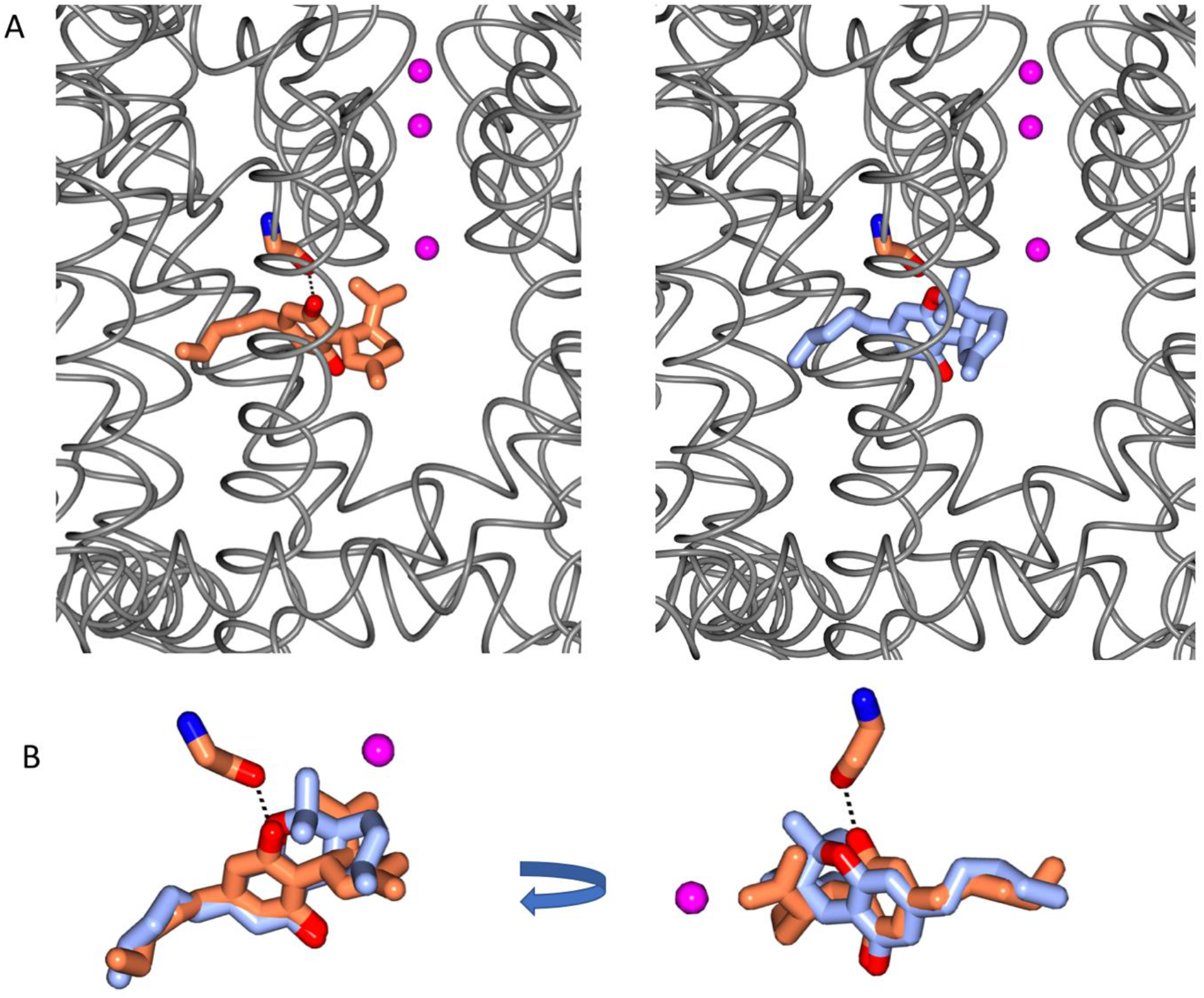
Comparison of CBD and THC in the CBD Binding Site. A) The location of the CBD site (left) and the modeled THC site (right). B) (left) The overlay of CBD (coral) and THC (blue) in the binding site shows the presence of the additional hydrogen bond between the protein and drug for CBD in comparison to that possible for THC. The view on the right side is rotated from the view on the left to visualise a different view of the alignment.

**Supplementary Fig. 8.**
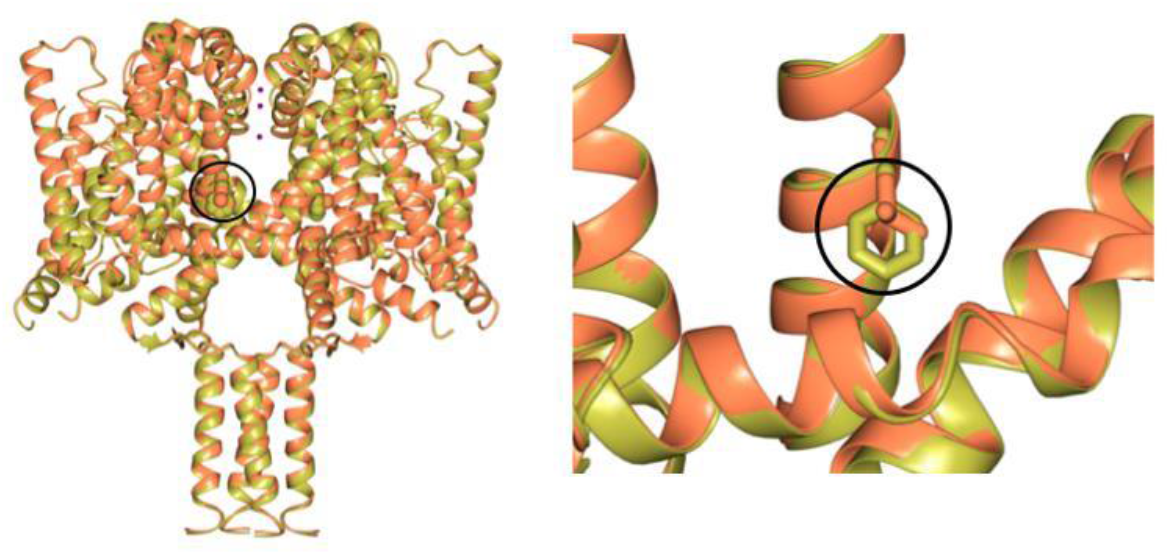
Comparisons of Wild Type NavMs (gold) and the NavMs_L_ Mutant (coral) structures. These figures show the (left) close overall similarities of the structures, and (right) in the detailed region surrounding mutated residue F208L (which is circled in both panels) (see Fig. 4 sequence).

**Supplementary Table 1.** Crystal Structure Parameters.

| *Data collection* | **NavMs<sub>L</sub> (PDB ID 6YZ2)** | **NavMs-CBD (PDB ID 6YZ0)** |
| --- | --- | --- |
| Wavelength (Å) | 0.97624 | 0.97966 |
| Space group | I422 | I422 |
| *Unit-cell parameters* |  |  |
| a, b, c (Å) | 108.94 ,108.94,209.37 | 108.4,108.4,208.2 |
| $\alpha, \beta, \gamma$ (°) | 90, 90, 90 | 90, 90, 90 |
| Resolution range (Å) | 104.68-2.20 (2.27-2.20) | 48.07-2.25(2.32-2.25) |
| Total number of observations | 687781 (56721) | 763702 (56037) |
| Total number unique | 32388 (2771) | 29849 (2708) |
| Completeness | 100.0 (100.0) | 100.0 (100.0) |
| Multiplicity | 21.2 (20.5) | 25.6 (20.7) |
| $\langle I/\sigma(I) \rangle$ | 17.4 (2.2) | 14.8 (1.5) |
| CC(1/2) | 0.999 (0.950) | 0.998 (0.861) |
| R<sub>merge</sub> all | 0.093 (1.20) | 0.123 (2.25) |
| R<sub>pim</sub> All | 0.021 (0.278) | 0.025 (0.520) |
| Solvent content (%) | 72.80 | 74.98 |
| Molecule per ASU | 1 | 1 |
| Wilson B factor (Å<sup>2</sup>) | 46.9 | 54.4 |
| *Refinement* |  |  |
| Resolution Range (Å) | 28.8-2.2 | 48.1-2.25 |
| R<sub>work</sub> | 0.24 | 0.23 |
| R<sub>free</sub> | 0.26 | 0.26 |
| Reflection, working | 32388 | 29848 |
| Reflection, free | 1619 | 1504 |
| Average B factor (all atoms) | 75.0 | 94.4 |
| RMS bond angle | 0.82 | 0.83 |
| RMS bond length (Å) | 0.008 | 0.008 |
| *Ramachandran Analysis* |  |  |
| Preferred region (%) | 96.6 | 94.8 |
| Allowed region (%) | 3.0 | 5.2 |
| Outliers (%) | 0.4 | 0.0 |

## References

Ahern CA, Payandeh J, Bosmans F, Chanda B. 2016. The hitchhiker’s guide to the voltage-gated sodium channel galaxy. J. Gen. Physiology 147:1–24.

Bagal SK, Marron BE, Owen RM, Storer RI, Swain NA. 2015. Voltage gated sodium channels as drug discovery targets. Channels 9:360–366.

Bagnéris C, DeCaen PG, Naylor CE, Pryde DC, Nobeli I, Clapham DE, Wallace BA. 2014. Prokaryotic NavMs channel as a structural and functional model for eukaryotic sodium channel antagonism. Proc. Natl. Acad. Sci. (USA) 111:8428–8433.

Catterall WA, Goldin AL, Waxman SG. 2005. International Union of Pharmacology XLVII. Nomenclature and structure-function relationships of voltage-gated sodium channels. Pharmacological Reviews 57:397–409.

Chen VB, Arendall III WB, Headd JJ, Keedy DA, Immormino RM, Kapral GJ, Murray LW, Richardson JS, Richardson DC. 2010. MolProbity: All-atom structure validation for macromolecular crystallography. Acta Cryst D66:12–21.

Chiu J, March PE, Lee R, Tillett D. 2004. Site-directed, ligase-independent mutagenesis (SLIM): a single-tube methodology approaching 100% efficiency in 4 h. Nucleic Acids Res. 32:e174.

Cross JH, Devinsky O, Marsh E, Miller I, Nabbout R, Scheffer IE, Thiele EA, Laux L, Wright S. 2017. Cannabidiol(CBD) reduces convulsive seizure frequency in Dravet syndrome: results of a multi-center, randomized, controlled trial (GWPCARE1) (CT. 001)

Devinsky O, Patel AD, Cross JH, Villanueva V, Wirrell EC, Privitera M, Greenwood SM, Roberts C, Checketts D, VanLandingham KE, Zuberi SM. 2018. Effect of cannabidiol on drop seizures in the lennox–gastaut syndrome. New England J. Medicine 378:1888–1897.

El-Din TMG, Lenaeus MJ, Zheng N, Catterall WA. 2018. Fenestrations control resting-state block of a voltage-gated sodium channel. Proc. Nat. Acad. Sci. (USA) 115:13111–13116.

Emsley P, Lohkamp B, Scott WG, Cowtan K. 2010. Features and development of Coot. Acta Cryst. D66:486–501.

Evans PR, Murshudov GN. 2013. How good are my data and what is the resolution? Acta Cryst. D69:1204–1214.

Ghovanloo MR, Shuart NG, Mezeyova J, Dean RA, Ruben PC, Goodchild SJ. 2018. Inhibitory effects of cannabidiol on voltage-dependent sodium currents. J. Biol. Chem. 293:16546–16558.

Hill AJ, Jones NA, Smith I, Hill CL, Williams CM, Stephens GJ, Whalley BJ. 2014. Voltage-gated sodium (NaV) channel blockade by plant cannabinoids does not confer anticonvulsant effects per se. Neuroscience Lett. 566:269–274.

Hille B. 1977. Local anesthetics: hydrophilic and hydrophobic pathways for the drug-receptor reaction. J. Gen. Physiology 69:497–515.

Humphrey W, Dalke A, Schulten K. 1996. VMD - Visual Molecular Dynamics. J. Mol. Graph. 14:33–38.

Jiang D, Shi H, Tonggu L, Gamal El-Din TM, Lenaeus MJ, Zhao Y, Yoshioka C, Zheng N, Catterall WA. 2019. Structure of the cardiacsodium channel. Cell 180:122–134.

Kabsch W. 2010 XDS. Acta Cryst. D66:125–132.

Kaplan DI, Isom, LL, Petrou S. 2016. Role of sodium channels in epilepsy. Cold Spring Harbor Persp. Med. 6:a022814.

Kaplan JS, Stella N, Catterall WA, Westenbroek RE. 2017. Cannabidiol attenuates seizures and social deficits in a mouse model of Dravet syndrome. Proc. Natl. Acad. Sci. (USA) 144:11229–11234.

Ke S, Ulmschneider MB, Wallace BA, Ulmschneider JP. 2018. Role of the interaction motif in maintaining the open gate of an open sodium channel. Biophysical J. 115:1920–1930.

Laskowski RA, MacArthur MW, Moss DS, Thornton JM. 1993. PROCHECK - a program to check the stereochemical quality of protein structures. J. App. Cryst. 26:283–291.

Marini C, Scheffer IE, Nabbout R, Suls A, De Jonghe P, Zara F, Guerrini R. 2011. The genetics of Dravet syndrome. Epilepsia 52:24–29.

Martin LJ, Corry B. 2014. Locating the route of entry and binding sites of benzocaine and phenytoin in a bacterial voltage gated sodium channel. PLoS Comp. Biol., 10:7.

Mason ER, Cummins TR. (2020) Differential Inhibition of human Nav1.2 resurgent and presistant sodium current by cannabidiol and GS967. Int. J. of Molecular Sciences 21, 2454.

McCoy AJ, Grosse-Kunstleve RW, Adams PD, Winn MD, Storoni LC, Read RJ. 2007. Phaser crystallographic software. J. App. Cryst. 404:658–674.

McNicholas S, Potterton E, Wilson KS, Noble MEM. 2011. Presenting your structures: the CCP4mg molecular-graphics software. Acta Cryst. D67:386–394.

Montini G, Booker J, Sula A, Wallace BA. 2018. Comparisons of voltage-gated sodium channel structures with open and closed gates and implications for state-dependent drug design. Biochem. Soc. Trans. 46:1567–1575.

Morelli MB, Amantini C, Liberati S, Santoni M, Nabissi M. 2013. TRP channels: new potential therapeutic approaches in CNS neuropathies. CNS & Neurological Disorders - Drug Targets 12: 274–293.

Murshudov G, Skub A, Lebede AA, Navraj S, Pannu RA, Steiner RA, Nicholls, Winn MD, Long F, Vagin AA. 2011. REFMACS for the refinement of macromolecular crystal structures. Acta Cryst. D67:355–367.

Naylor CE, Bagnéris C, DeCaen PG, Sula A, Scaglione A, Clapham DE, Wallace BA. 2016. Molecular basis of ion permeability in a voltage-gated sodium channel. EMBO J. 35:820–830.

Ouyang W, Jih TY, Zhang TT, Correa AM, Hemmings HC. 2007. Isoflurane inhibits NaChBac, a prokaryotic voltage-gated sodium channel. J. Pharm. Exptl. Therapeutics 322: 1076–1083.

Patel RR, Barbosa C, Brustovetsky T, Brustovetsky N, Cummins TR. 2016. Aberrant epilepsy-associated mutant Nav1.6 sodium channel activity can be targeted with cannabidiol. Brain 139:2164–2181.

Payandeh J, Scheuer T, Zheng N, Catterall WA. 2011. The crystal structure of a voltage-gated sodium channel. Nature 475:353–358.

Pisanti S, Malfitano AM, Ciaglia E, Lamberti A, Ranieri R, Cuomo G, Abate M et al. 2017. Cannabidiol: State of the art and new challenges for therapeutic applications. Pharm. Therapeutics 175:133–115.

Pumroy RA et al. 2019. Molecular mechanism of TRPV2 channel modulation by cannabidiol. Elife 8:48792.

Qin N, Neeper MP, Liu Y, Hutchinson TL, Lubin ML, Flores CM. 2008. TRPV2 is activated by cannabidiol and mediates CGRP release in cultured rat dorsal root ganglion neurons. J. Neuroscience 28:6231–6238.

Rosenberg EC, Tsien RW, Whalley BJ, Devinsky O. 2015. Cannabinoids and epilepsy. Neurotherapeutics 12: 747–768.

Rosenberg EC., Patra PH, Whalley BJ 2017. Therapeutic effects of cannabinoids in animal models of seizures, epilepsy, epileptogenesis, and epilepsy-related neuroprotection. Epilepsy & Behavior 70:319–327.

Sarker SD, Nahar L. 2020. Cannabidiol (CBD) – An update. Trends Phytochemical Res. 4:1–2.

Sievers FA, Wilm D, Dineen TJ, Gibson K, Karplus W, Li R, Lopez H, McWilliam M, Remmert J, Söding JD Thompson, Higgins GD. 2011. Fast, scalable generation of high-quality protein multiple sequence alignments using Clustal Omega. Mol. Systems Biol. 7: 539.

Smart OS, Goodfellow JM, Wallace BA. 1993. The pore dimensions of gramicidin A. Biophys J. 65:2455–2460.

Sula A, Wallace BA. 2017. Interpreting the functional role of a novel interaction motif in prokaryotic sodium channels. J. Gen. Physiol. 149:613–622.

Sula A, Booker J, Ng LCT, Naylor CE, DeCaen PG, Wallace BA. 2017. The complete structure of an activated open sodium channel. Nature Comms. 8:14205.

Ulmschneider MB, Bagnéris C, McCusker EC, DeCaen PG, Delling M, Clapham DE, Ulmschneider JP, Wallace BA. 2013. Molecular dynamics of ion transport through the open conformation of a bacterial voltage-gated sodium channel. Proc. Natl. Acad. Sci. (USA) 110:6364–6369.

Waterhouse AM, Procter JB, Martin DMA, Clamp M, Barton GJ. 2009. Jalview Version 2 – a multiple sequence alinment editor and analysis workbench. Bioinformatics. 25 (9): 1189–1191.

Watkins AR. 2019. Cannabinoid interactions with ion channels and receptors. Channels 13:162–167.

Winn MD et al. 2011. Overview of the CCP4 suite and current developments. Acta Cryst. D67:235–242.

